# Biting and resting preferences of malaria vectors in The Gambia

**DOI:** 10.1101/2020.10.08.331165

**Authors:** Majidah Hamid-Adiamoh, Davis Nwakanma, Benoit Sessinou Assogba, Mamadou Ousmane Ndiath, Umberto D’Alessandro, Yaw A. Afrane, Alfred Amambua-Ngwa

## Abstract

**Background:** The scale-up of indoor residual spraying and long-lasting insecticidal nets, together with other interventions have considerably reduced the malaria burden in The Gambia. This study examined the biting and resting preferences of the local insecticide-resistant vector populations few years following scale-up of anti-vector interventions.

**Method:** Indoor and outdoor-resting *Anopheles gambiae* mosquitoes were collected between July and October 2019 from ten villages in five regions in The Gambia using pyrethrum spray collection (indoor) and prokopack aspirator from pit traps (outdoor). Polymerase chain reaction assays were performed to identify molecular species, insecticide resistance mutations, *Plasmodium* infection rate and host blood meal.

**Results:** A total of 844 mosquitoes were collected both indoors (421, 49.9%) and outdoors (423, 50.1%). Four main vector species were identified, including *An. arabiensis* (indoor: 15%, outdoor: 26%); *An. coluzzii* (indoor: 19%, outdoor: 6%), *An. gambiae s*.*s*. (indoor: 11%, outdoor: 16%), *An. melas* (indoor: 2%, outdoor: 0.1%) and hybrids of *An. coluzzii-An. gambiae* (indoors: 3%, outdoors: 2%). A significant preference for outdoor resting was observed in *An. arabiensis* (Pearson *X*^*2*^=22.7, df=4, P<0.001) and for indoor resting in *An. coluzzii* (Pearson *X*^*2*^=55.0, df=4, P<0.001). Prevalence of the voltage-gated sodium channel (*Vgsc*)*-1014S* was higher in the indoor-resting (allele freq. = 0.96, 95%CI: 0.78–1) than outdoor-resting (allele freq. = 0.82, 95%CI: 0.76–0.87) *An. arabiensis* population. For *An. coluzzii*, the prevalence of most mutation markers were higher in the outdoor (allele freq. = 0.92, 95%CI: 0.81–0.98) than indoor-resting (allele freq. = 0.78, 95%CI: 0.56–0.86) mosquitoes. Sporozoite positivity rate was 1.3% (95% CI: 0.5–2%). Indoor-resting *An. coluzzii* had mainly fed on human blood while indoor-resting *An. arabiensis*, animal blood.

**Conclusion:** The indoor-resting behavior of *An. arabiensis* that preferred animal blood and had low sporozoite rates, may be determined by the *Vgsc-1014S* mutation. Control interventions may include complementary vector control approaches such as zooprophylaxis.

## Introduction

Successful implementation of indoor residual spraying (IRS) and long-lasting insecticidal nets (LLINs) has hugely contributed to the malaria decline observed in sub-Saharan Africa [1]. These interventions reduce transmission by primarily limiting human contact with human-feeding (anthropophagic), indoor-feeding (endophagic) and indoor-resting (endophilic) vectors [2]. Unfortunately, these measures also induce selection for physiological and behavioral resistance in vector populations, resulting in reduced mosquito susceptibility to most of the current insecticides used for LLINs and IRS [3], as well as increased exophilic behavioral phenotypes in primarily endophilic vectors [4]. Moreover, residual transmission where LLINs and IRS use is extensive, is maintained by vectors with physiological and behavioral resistance [5]. Therefore, studying the behavioral dynamics of vector populations during the scale up of vector control interventions will assist in determining the appropriate response to emerging behavioral changes.

Malaria burden in The Gambia has declined significantly over the last decades with vector control approaches being a major component of intervention, coordinated and implemented by The Gambia National Malaria Control Program (GNMCP). Following the World Health Organization (WHO) Global Plan for Insecticide Resistance Management (GPIRM), the GNMCP has consistently implemented rotational use of different classes of insecticides for IRS, to curtail dichlorodiphenyltrichloroethane (DDT) and deltamethrin resistance. For IRS, DDT was replaced initially by deltamethrin and bendiocarb, and since 2017 by pirimiphos-methyl (actellic 300CS) [6]. Similarly, LLINs intervention has been stable over the years and Gambia has recorded successful LLINs coverage as high as 90% [7,8].

Despite such successes, residual transmission has become increasingly spatially heterogeneous, with its intensity increasing from western to eastern Gambia, and could have been driven by specific vector population dynamics [9]. The major vector species, namely *Anopheles arabiensis, An. coluzzii* and *An. gambiae sensu stricto* (*s*.*s*.) are variably distributed throughout the country. *An. arabiensis* is most prevalent in the eastern Gambia while *An. coluzzii* and *An. gambiae s*.*s*. inhabit the western region [10,11]. However, *An. arabiensis* has been recently found throughout the country [12], indicating possible replacement due to successful control of other sibling species [13,14]. Moreover, the population prevalence of each vector species varies by season, whereby *An. arabiensis* and *An. coluzzii* are dominant throughout the rainy season, while *An. gambiae s*.*s*. become rarest early in the onset of dry season [10,11]. DDT and pyrethroid resistance has been reported at various degrees in all vectors, that continue to be highly susceptible to carbamates and organophosphates [12,15,16].

Host seeking and resting behavior of vectors are important metrics to evaluate the impact of control and resistance management strategies [17]. Vector behavioral adaptation, resistance selection and persistent transmission could increase during extensive scale-up of interventions, and this information can only be captured by real-time surveillance [18,19]. Hence, national malaria control programs should actively monitor behavioral dynamics in the local vector population, to inform decisions.

In The Gambia, DDT and pyrethroid resistance is widespread and associated with residual transmission [12,15]. However, the effect of control activities on vectors feeding and resting behavior remains unclear. The biting and resting preferences of *An. gambiae sensu lato* (*s*.*l*.) populations was investigated in The Gambia following few years of intensive vector control interventions.

## Materials and methods

### *Anopheles gambiae s*.*l*. collection

Indoor and outdoor-resting adult mosquitoes were sampled from July to October 2019, during the malaria transmission season across five administrative regions in The Gambia, namely Central River Region (CRR), Lower River Region (LRR), North Bank Region (NBR), Upper River Region (URR) and West Coast Region (WCR). WCR is a coastal area characterized by mangrove swamps. The remaining regions are mainly inland and have forest vegetation. Rice is mainly cultivated in CRR while cereals farming is common in all regions. Two villages were selected from each region and most of the villages are GNMCP surveillance sites with high LLIN and IRS coverage. Malaria transmission is highest in URR compared to other regions in The Gambia [7].

Indoor-resting mosquitoes were collected from sleeping rooms using pyrethrum spray collection (PSC). Twenty houses per village, at least 50m apart from each other, were randomly selected. In each village, collections were done for two consecutive days, with ten houses sampled per day. Outdoor-resting mosquitoes were sampled from pit shelter traps using prokopak aspirator. Three pit shelter traps that were 10m away from the selected compounds, were placed at different parts in each village. Both indoor and outdoor collections were conducted from 06.00 am to 09.00am in every collection day.

### Mosquito identification

Morphological identification of female *An. gambiae s*.*l*. was done using identification keys as described by Gillies & Coetzee [20]. Afterwards, mosquitoes were stored individually in 96% ethanol in 1.5ml Eppendorf tube until DNA extraction. DNA was extracted separately from abdomen and head/thoraces of individual mosquitoes using Qiagen QIAxtractor robot. Species-specific genotyping PCR to identify *An. arabiensis, An. melas* and *An. gambiae* was performed as previously described [21]. This was followed by restriction enzyme digestion to specifically identify *An. coluzzii, An. gambiae s*.*s*. and their hybrids (*An. coluzzii*-*An. gambiae s*.*s*.) [22].

### Insecticide resistance markers identification

Screening for molecular markers of target-site resistance to carbamates, DDT, pyrethroids and organophosphates was done on all samples using a probe-based assay (TaqMan SNP genotyping) [23]. The following markers were investigated: voltage-gated sodium (*Vgsc*)*-1014F, Vgsc-1014S* and *Vgsc-1575Y* associated with target-site mutation to DDT and pyrethroids [24–26]. Acetylcholine esterase (*Ace*)*-119S*, marker for carbamate and organophosphate resistance [27] and glutathione-S-transferase epsilon 2 (*Gste2*)*-114T*, involved in metabolic resistance to DDT [28] were also assayed. The TaqMan allelic discrimination assay is a multiplex real time PCR with primers and probes specific for each insecticide target gene and discriminate susceptible (wild type) and resistant (mutant) alleles based on probe fluorescence signals [29].

### Plasmodium sporozoite detection

DNA extracted from mosquito head and thoraces was used to detect sporozoites of *Plasmodium falciparum, P. ovale, P. malariae* and *P. vivax* species, employing TaqMan SNP genotyping protocol [30] which enables discriminatory identification of circum-sporozoites (CSPs) of *P. falciparum* from *P. ovale, P. malariae* and *P. vivax* CSPs. Genomic DNA specific to each of these *Plasmodium* species were analyzed in each assay as positive controls.

### Blood meal identification

Extracted DNA from engorged mosquito abdomens were amplified using modified multiplex PCRs with primers targeting cytochrome B genes of human and animal hosts including chicken, cow, dog, donkey, goat, horse and pig [31,32].

## Statistical analyses

The proportion of each mosquito species in relation to the total number of mosquitoes captured from each region was calculated in percentage, as well as allele frequencies of indoor and outdoor-resting mosquitoes. Sporozoite positivity rate was the proportion of PCR positive mosquitoes among all mosquitoes tested. Human (HBI) and animal blood meal indices were estimated as the proportion of mosquitoes positive for human or animal hosts among those positive for all hosts. Mean differences between HBI and animal blood meal indices by vector species and resting locations were analyzed by ANOVA. Statistical analyses were done using Stata/IC 15.0 (2017 StataCorp LP).

## Results

### Anopheles species distribution and their resting behavior

A total of 844 *An. gambiae s*.*l*. mosquitoes were collected from the five regions. Four main vector species were identified, namely *An. arabiensis* (N=350, 41%); *An. coluzzii* (N=214, 25%), *An. gambiae s*.*s*. (N=224, 27%) and *An. melas* (N=17, 2%). Hybrids of *An. coluzzii-An. gambiae s*.*s*. were also detected (N=39, 5%). Most mosquitoes were collected from URR (642, 76%), followed by LRR (97, 11%) and then the other regions (Fig 1).

**Fig 1:**
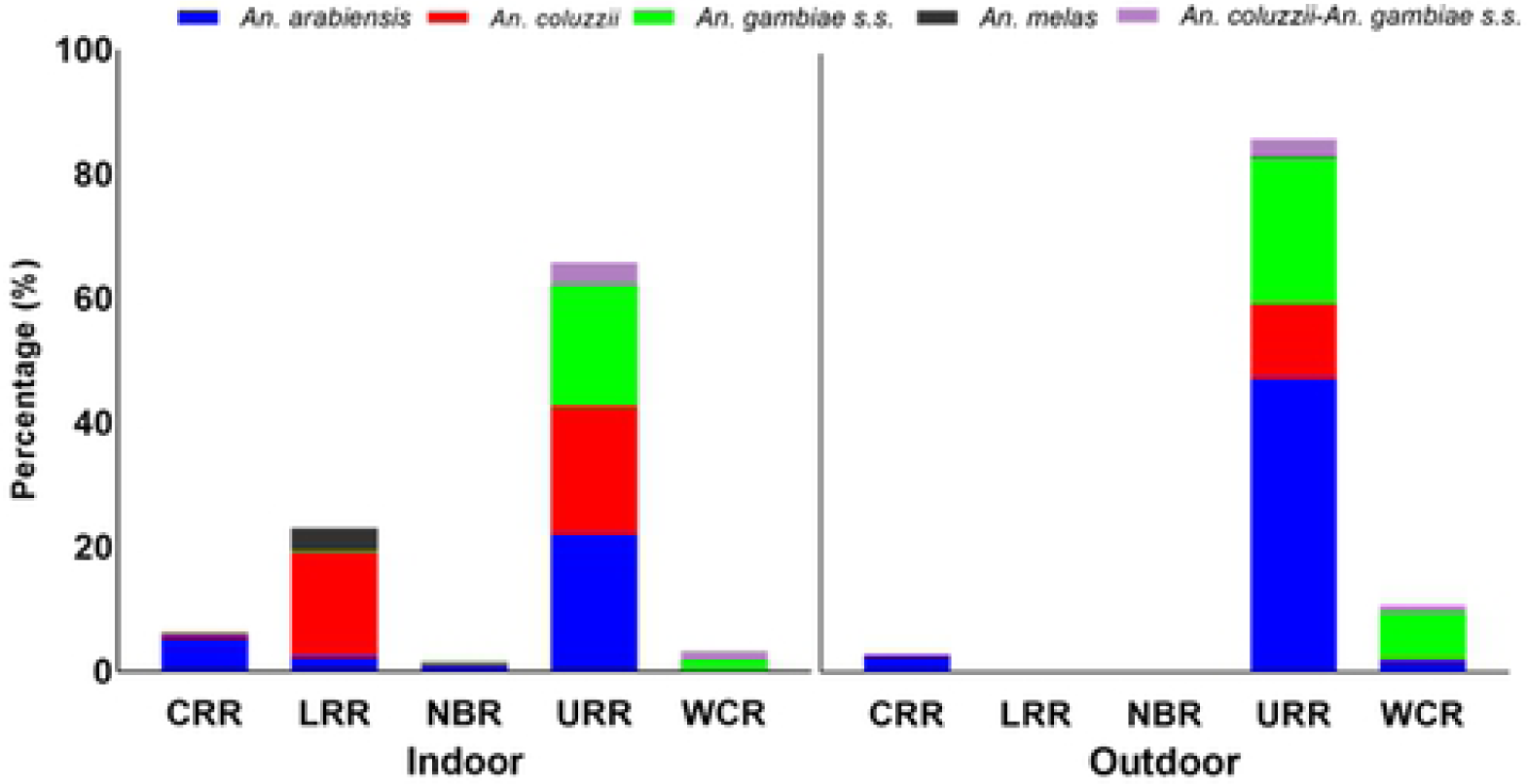
Distribution of *Anopheles gambiae s*.*l* by region as collected indoors and outdoors. *An. col*uzzii-An. *gambiae s*.*s*. are the hybrids of *An. coluzzii* and *An. gambiae s*.*s*. Mosquitoes were collected from 5 regions: CRR-central river region, LRR-lower river region. NBR-north bank region. URR-upper river region and WCR-West coast region.

Overall, the number of mosquitoes resting indoors (421, 49.9%) and outdoors (423, 50.1%) were similar. Nevertheless, the resting preference varied by species. A significantly higher proportion of *An. arabiensis* were found outdoor (26.1%) than indoor (15.4%) (Pearson *X*^*2*^=22.7, df=4, P<0.001) while both *An. coluzzii* (19.1% indoor and 6.3% outdoor, Pearson *X*^*2*^=55.0, df=4, P<0.001) and *An. melas* (1.9% indoor and 0.1% outdoor, Pearson *X*^*2*^=13.3, df=4, P<0.01) preferred resting indoor. For *An. gambiae s*.*s*. (10.9% indoor and 15.6% outdoor, Pearson *X*^*2*^=7.0, df=4, P<0.1) and *An. coluzzii-An. gambiae s*.*s*. hybrids (2.6% indoor and 2% outdoor, Pearson *X*^*2*^=0.7, df=4, P<0.95), there was no significance difference between resting indoor and outdoor. In URR, the region with the highest malaria transmission in The Gambia, *An. arabiensis* was most abundant vector (45.8%, 294) (indoor: 14.5%, outdoor: 31.3%), followed by *An. gambiae s*.*s*. (28.4%, 182) (indoor: 12.8%, outdoor: 15.6%) and *An. coluzzii* (21.5%, 138) (indoor: 13.6%, outdoor: 7.9%). No *An. gambiae s*.*s*. was collected in CRR while *An. melas* was mainly found in LRR (N=15). All mosquitoes collected from LRR and NBR were resting indoors. The hybrids of *An. coluzzii* and *An. gambiae s*.*s*. were mainly found in URR (indoor: 2.3%, outdoor: 1.9%) and WCR (indoor: 10%, outdoor: 8.3%).

### Distribution of voltage-gated sodium channel (Vgsc) mutation markers in the vectors

Vgsc point mutations associated with DDT and pyrethroid resistance were highly prevalent and detected at varying frequencies in all vector species across all regions. Overall, *An. arabiensis* was found resting indoors when resistance allele frequency was higher in the indoor population, whereas *An. coluzzii* were resting outdoors with higher outdoor resistance. No consistent resting preference was observed in *An. gambiae* in the presence of mutations.

*Vgsc-1014S* mutation was found predominantly in indoor-resting vector populations (Table 1). In *An. arabiensis*, the mutation was more frequent in the indoor-resting than outdoor-resting mosquitoes regardless of the region. V*gsc-1014S* was also the only mutation identified in *An. gambiae s*.*s*. and *An. melas* when found resting indoors.

**Table 1:** Frequencies of insecticide resistance alleles on VGSC, GST and AChE loci in *Anopheles gambiae s*.*l*. populations from all study regions.

| Region | Anopheles species | *Vgsc-1014F* |  | *Vgsc-1014S* |  | *Vgsc-1575Y* |  | *GSTe2-114T* |  | *Ace1-119S* |  |
| --- | --- | --- | --- | --- | --- | --- | --- | --- | --- | --- | --- |
|  |  | Indoor | Outdoor | Indoor | Outdoor | Indoor | Outdoor | Indoor | Outdoor | Indoor | Outdoor |
| URR | *An. arabiensis* (N=294) | 0.05 | 0.02 | 0.91 | 0.82 | 0 | 0.004 | 0 | 0.01 | 0 | 0 |
|  | *An. coluzzii* (N=138) | 0.74 | 0.92 | 0.25 | 0.04 | 0.68 | 0.9 | 0.78 | 0.9 | 0 | 0 |
|  | *An. gambiae* ss (N=182) | 1 | 0.99 | 0 | 0.01 | 0.96 | 0.98 | 0.98 | 0.99 | 0.05 | 0.04 |
|  | *An. col./ gam* (N=27) | 0.93 | 1 | 0 | 0 | 0.87 | 1 | 0.87 | 0 | 0.07 | 0 |
| LRR | *An. arabiensis* (N=10) | 0 | - | 0.9 | - | 0 | - | 0.1 | - | 0 | - |
|  | *An. coluzzii* (N=71) | 0.66 | - | 0.3 | - | 0 | - | 0 | - | 0 | - |
|  | *An. gambiae* ss (N=1) | 0 | - | 1 | - | 0 | - | 0 | - | 0 | - |
|  | *An. melas* (N=15) | 0 | - | 1 | - | 0 | - | 0 | - | 0 | - |
| WCR | *An. arabiensis* (N=8) | - | 0.13 | - | 0.88 | - | 0 | - | 0 | - | 0 |
|  | *An. coluzzii* (N=1) | - | 1 | - | 0 | - | 0 | - | 0 | - | 0 |
|  | *An. gambiae* ss (N=40) | 0.25 | 0.13 | 0.75 | 0.84 | 0.13 | 0.06 | 0 | 0 | 0 | 0 |
|  | *An. col./ gam* (N=11) | 0 | 0 | 1 | 1 | 0 | 0 | 0.17 | 0 | 0 | - |
| CRR | *An. arabiensis* (N=34) | 0 | 1 | 0.96 | 0 | 0 | 0 | 0.1 | 0.27 | 0 | 0 |
|  | *An. coluzzii* (N=3) | - | 1 | - | 0 | 0 | 0 | 0 | 0 | 0 | 0 |
|  | *An. melas* (N=1) | 0 | - | 1 | - | 0 | - | 0 | - | - | 0 |

Vgsc- voltage-gated sodium channel. GSTe2-glutathione-s-transferase epsilon 2. Ace1-Acetylcholine esterase1.

In *An. arabensis* resting indoors in URR, *Vgsc-1014S* frequency was higher in the indoor-(allele freq. = 0.91, 95%CI: 0.84–0.96) than outdoor-resting (allele freq. = 0.82, 95%CI: 0.76–0.87) mosquitoes. Moreover, *Vgsc-1014S* was the only mutation identified in this species when found resting indoors (allele freq. = 0.96, 95%CI: 0.78–1) in CRR. Similarly in URR, the *Vgsc-1014S* mutation in *An. coluzzii* was higher in the indoor (allele freq. = 0.25, 95%CI: 0.17–0.36) than outdoor-resting mosquitoes (allele freq. = 0.04, 95%CI: 0.005–1.3). In LRR, the mutation was found only in indoor-resting mosquitoes (allele freq. = 0.3, 95%CI: 0.19–0.42). Conversely in WCR, the mutation was common in *An. gambiae s*.*s*. and higher among outdoor-(allele freq. = 0.84, 95%CI: 0.67–0.95) than indoor-resting (allele freq. = 0.75, 95%CI: 0.35–0.97) mosquitoes.

*Vgsc-1014F* was almost fixed in most mosquitoes, except *An. arabiensis*. It was also more common in the outdoor-than indoor-resting mosquitoes. More specifically in URR, the mutation was found higher in outdoor-resting (allele freq. = 0.92, 95%CI: 0.81–0.98) than the indoor-resting *An. coluzzi* population (allele freq. = 0.74, 95%CI: 0.63–0.82). Likewise, in the hybrid population of *An. coluzzi* and *An. gambiae s*.*s*., the mutation was fixed and higher in the outdoor-resting (allele freq. = 1, 95%CI: 0.74–1) than indoor-resting (allele freq. = 0.93, 95%CI: 0.80–1) mosquitoes. The mutation was similarly fixed in both the indoor (allele freq. = 1, 95%CI: 0.96–1) and outdoor (allele freq. = 0.99, 95%CI: 0.95–1) *An. gambiae s*.*s*. populations. Conversely in WCR, *Vgsc-1014F* was more frequent in *An. gambiae s*.*s*. resting indoors (allele freq. = 0.25, 95%CI: 0.03– 0.65) than the outdoor population (allele freq. = 0.13, 95%CI: 0.04–0.29). Whereas in LRR, where only mosquitoes resting indoors were caught, this mutation was most common in *An. coluzzii* (allele freq. = 0.66, 95%CI: 0.81–0.98).

*Vgsc-1575Y* and *GSTe2-114T* were found mostly in URR and were more frequent in outdoor-resting mosquitoes. The mutations were almost fixed in *An. gambiae s*.*s*. regardless of resting place (allele freq. = 0.96-1, 95% CI: 0.92–1.2). When *An. coluzzii* was found resting outdoors also in this region, these mutations were higher (allele freq, = 0.9, 95% CI: 0.79–0.97) than in their indoor-resting counterpart (allele freq, = 0.68-0.78, 95% CI: 0.56–0.86). The hybrids of *An. coluzzii* and *An. gambiae s*.*s*. with higher and fixed *Vgsc-1575Y* mutation were equally resting outdoors (allele freq. = 1, 95% CI: 0.74–1) while those found resting indoors were carrying only the *GSTe2-114T* mutation (allele freq. = 0.87, 95% CI: 0.60–0.98).

The carbamate and organophosphate resistance marker, acetylcholine esterase (*Ace*)*-119S* was detected only in 8 (4 indoor and 4 outdoor) *An. gambiae s*.*s*. and in one hybrid specimen in URR.

### Sporozoite infection rate

*Plasmodium falciparum* sporozoites were detected in 11 out of 844 mosquitoes (Table 2), representing a 1.3% (95% CI: 0.5–2%) infection rate. All the infected mosquitoes were caught in URR, of which six were resting indoors and five resting outdoors. Outdoor-resting *An. arabiensis* were mostly infected (36%, 4/11), followed by indoor-resting *An. gambiae s*.*s*. (27%, 3/11) and *An. arabiensis* (18%, 2/11). One each of outdoor-resting *An. coluzzii* and *An. coluzzii*-A*n. gambiae s*.*s*. hybrid were also infected.

**Table 2:** Sporozoite positivity rate in the eleven vector species that were infected based on their resting locations.

|  | *An. arabiensis*<br>proportion (n) | *An. coluzzii*<br>proportion (n) | *An. gambiae* s.s.<br>proportion (n) | *An. coluzzii-An.*<br>*gambiae* s.s.<br>proportion (n) |
| --- | --- | --- | --- | --- |
| Indoor | 0.18 (2) | 0.09 (1) | 0.27 (3) | 0 |
| Outdoor | 0.36 (4) | 0 | 0 | 0.09 (1) |

n= number of mosquitoes positive for sporozoite detection. Proportion = the number positive per species divided by overall positive (11).

### Host blood meal preference

Host blood meal origin was determined in 251 randomly selected engorged mosquito abdomens. Overall, animal and human blood meal indices were higher for indoor-than outdoor-resting mosquitoes (Table 3). In all vector species, most blood meal (91%) had animal origin. Indoor-resting *An. coluzzii* had the highest preference for human blood while indoor-resting *An. arabiensis* had most preference for animal blood.

**Table 3:** Human and animal blood meal preferences of the indoor and outdoor-resting vector species in combined study sites.

|  | *An. arabiensis* |  | *An. coluzzii* |  | *An. gambiae ss* |  | *An. col./An. gambiae ss* |  |
| --- | --- | --- | --- | --- | --- | --- | --- | --- |
|  | Indoor (n) | Outdoor(n) | Indoor(n) | Outdoor(n) | Indoor(n) | Outdoor(n) | Indoor(n) | Outdoor(n) |
| Human | 0.01 (3) | 0.004 (1) | 0.03 (7) | 0 | 0.01 (2) | 0.01 (2) | 0 | 0.004 (1) |
| Animal | 0.23 (58) | 0.16 (40) | 0.12 (30) | 0.09 (23) | 0.12 (29) | 0.13 (33) | 0.04 (9) | 0.02 (6) |
| Human + Animal | 0.004 (1) | 0.004 (1) | 0.01 (3) | 0 | 0.01 (2) | 0 | 0 | 0 |
| HBI | 2 | 0.8 | 4 | 0 | 1 | 0.8 | 0 | 0.4 |
| Animal blood indices | 23 | 16 | 12 | 9 | 12 | 13 | 4 | 2 |

Proportion (number). HBI= Human blood index.

## Discussion

Insecticide resistance is currently widespread among malaria vectors in The Gambia [12,15], resulting in insecticide rotation for IRS and more recently the use of actellic, an organophosphate insecticide, as recommended by WHO [33]. It is unclear how such vector control interventions have influenced the vectors’ feeding and resting behavior, and malaria transmission dynamics. In this study, *An. arabiensis* had a marked preference for outdoor resting and *An. coluzzii* and *An. melas* for indoor resting. Resting location was similar for *An. gambiae s*.*s*. and *An. coluzzii-An. gambiae s*.*s*. hybrid populations. Moreover, local vectors had a marked preference for animal blood. The sporozoite infection rate was low and infectious mosquitoes were mainly outdoor-resting *An*. a*rabiensis*.

*An. arabiensis* tended to rest indoors if they had resistance mutations. This was particularly evident for the *Vgsc-1014S* which seems to influence the resting behavior of this vector species. Vectors with this mutation, which is not fixed yet, may prefer to rest indoors as it protects against the effect of IRS and LLINs [37]. It would be worthwhile to further explore how this mutation modulates resting behavior in vector species. Conversely, *Vgsc-1014F* seems to be fixed in most vector populations in West Africa [34,35], and this may explain why vectors with this mutation do not have a specific resting behavior [36].

*An. coluzzii* displayed an outdoor-resting behavior when genotypically more resistant than the indoor population. This was consistent across all mutation markers except *Vgsc-1014S;* and was observed mainly in URR, where most mosquitoes were caught. The association between genotypic resistance and outdoor-resting behavior in *An. coluzzii* was recently reported in Northern Ghana [36]. However, for *An. gambiae s*.*s*., the mutation frequency did not vary by resting location. *Vgsc-1014F* was the main mutation in *An. gambiae s*.*s* at fixed frequencies and did not have any significant effect on resting behavior.

The vector populations analyzed had a higher preference for animal than human blood meal and the overall sporozoite rate was also low. Given the current low malaria prevalence in The Gambia [9], a low sporozote rate is expected. These may reflect the impact of the scaled-up in IRS and LLINs program in the study sites which seems to successfully limit mosquito access to human blood meal indoors and consequently reducing transmission, as previously reported [39,40]. The observed choice of animal blood by majority of the vectors could lead to increase in vector population that may eventually resort to biting humans in the long run and become difficult to control.

Furthermore, the proportion of vectors resting indoors and that have taken a blood meal either from human and animal source, could be a concern for the effectiveness of vector control measures. This shows that the vectors took blood meal from animals outdoors and later went indoors to rest regardless of the presence of IRS and LLINs, indicating that these vectors are resistant to the insecticides being used. This behavior was as previously demonstrated where blood-fed mosquitoes were more resistant than their unfed counterpart [41]. Notably, alternative vector control methods such as treatment of animals with endectocides [42] and zooprophylaxis [43], could be promising tools that could be adopted by the Gambia National Malaria Control Program.

The significant preference for outdoor resting by *An. arabiensis* and indoor resting in *An. coluzzii* prevalent in URR may explain the high intensity of malaria transmission in this region [7,9], which may be driven by these vectors. *An. coluzzii* is highly anthropophagic, endophilic and an efficient vector of malaria (44), traits that facilitate its contact with human indoors as observed here. A previous study in this setting corroborated the finding here where HBI as high as 80% was documented in *An. coluzzii* and *An. gambiae s*.*s*. [10]. Further, a relatively high vector parity rate was recently reported in the same vector species [9]. Moreover, *An. arabiensis* is known for its exophilic behavior that increases outdoor transmission in unprotected humans outside LLINs [45].

The composition of the vector species was consistent with previous studies in the Gambia where the most abundant vector was *An. arabiensis*, followed by *An. gambiae s*.*s*. and, *An. coluzzii* along with their hybrids [12,15,16]. Low density of *An. melas* found was as a result of our choice of villages in the West, which were not located in the coastal regions where this species breeds in salty water [46–48]. Remarkably, predominance of *An. arabiensis* could be as a result of its outdoor-resting preference to avoid insecticide used in IRS and LLINs [49]. This leaves the highly anthropophilic and endophilic species more exposed to vector interventions, possibly leading to relative advantage that maintains the exophilic population and malaria transmission [45].

## Conclusion

The study observed an indoor-resting behavior in *An. arabiensis* that were carrying *Vgsc-1014S* mutation and outdoor-resting behavior in *An. coluzzii* populations having other mutations. However, preference for outdoor resting was predominant in *An. arabiensis* and indoor resting in *An. coluzzii* populations. No specific preference for indoor or outdoor-resting behavior was demonstrated in *An. gambiae s*.*s*. and remaining vector species. An overall high preference for animal blood meal was found in the vector populations. Low rate of mosquito infectivity was identified likely due to high coverage of LLINs and IRS in the study regions. As malaria transmission remains low in The Gambia, which is in earnest preparation for pre-elimination phase, the magnitude of genotypic resistance observed in this study suggests a serious threat to the success of vector intervention in pre-elimination programs. Finally, the observed preference for animal host by vector populations, recommends the consideration of veterinary endectocides and zooprophylaxis as complementary vector control measures.

## Acknowledgements

We thank Messrs Musa Jawara, and Mamlie Touray for their assistance in the field work for this study.

## References

1. WHO. World Malaria Report 2019. Geneva. World Health Organization; 2019.

2. Killeen GF, Smith TA. Exploring the contributions of bed nets, cattle, insecticides and excitorepellency to malaria control: a deterministic model of mosquito host-seeking behavior and mortality. Trans R Soc Trop Med Hyg. 2007;9:867–80.

3. Ranson H, Guessan RN, Lines J, Moiroux N, Nkuni Z, Corbel V. Pyrethroid resistance in African anopheline mosquitoes: what are the implications for malaria control? Trends Parasitol 2011;27:91–8.

4. Gatton ML, Chitnis N, Churcher T, Donnelly MJ, Ghani AC, Charles HJ, et al. The importance of mosquito behavioral adaptations to malaria control in Africa. Evolution. 2013;1218–30.

5. Killeen GF. Characterizing, controlling and eliminating residual malaria transmission. Malar J. 2014;13:330.

6. Gambia National Malaria Control Program (GNMCP). Malaria Indicator Survey, Banjul. 2011.

7. Mwesigwa J, Okebe J, Affara M, Di Tanna GL, Nwakanma D, Janha O, et al. On-going malaria transmission in The Gambia despite high coverage of control interventions: A nationwide cross-sectional survey. Malar J. 2015;14:1–9.

8. Pinder M, Jawara M, Jarju LBS, Salami K, Jeffries D, Adiamoh M, et al. Efficacy of indoor residual spraying with dichlorodiphenyltrichloroethane against malaria in Gambian communities with high usage of long-lasting insecticidal mosquito nets: A cluster-randomised controlled trial. Lancet. 2015;385(9976):1436–46.

9. Mwesigwa J, Achan J, Di Tanna GL, Affara M, Jawara M, Worwui A, et al. Residual malaria transmission dynamics varies across The Gambia despite high coverage of control interventions. PLoS One. 2017;12:1–24.

10. Caputo B, Nwakanma D, Jawara M, Adiamoh M, Dia I, Konate L, et al. Anopheles gambiae complex along the Gambia river, with particular reference to the molecular forms of An. gambiae s.s. Malar J. 2008;7:182.

11. Jawara M, Pinder M, Drakeley CJ, Nwakanma DC, Jallow E, Bogh C, et al. Dry season ecology of Anopheles gambiae complex mosquitoes in the Gambia. Malar J. 2008;7:156.

12. Opondo KO, Jawara M, Cham S, Jatta E, Jarju L, Camara M, et al. Status of insecticide resistance in Anopheles gambiae (s.l.) of the Gambia. Parasit Vectors. 2019;12:1–287.

13. Kitau J, Oxborough RM, Tungu PK, Matowo J, Malima RC, Magesa SM, et al. Species shifts in the Anopheles gambiae complex: Do LLINs successfully control Anopheles arabiensis? PLoS One. 2012; 7:e31481.

14. Sougoufara S, Harry M, Doucouré S, Sembène PM, Sokhna C. Shift in species composition in the Anopheles gambiae complex after implementation of long-lasting insecticidal nets in Dielmo, Senegal. Med Vet Entomol. 2016;30:365–8.

15. Opondo KO, Weetman D, Jawara M, Diatta M, Fofana A, Crombe F, et al. Does insecticide resistance contribute to heterogeneities in malaria transmission in the Gambia? Malar J. 2016;15:1–10.

16. Wilson AL, Pinder M, Bradley J, Donnelly MJ, Hamid-Adiamoh M, Jarju LBS, et al. Emergence of knock-down resistance in the Anopheles gambiae complex in the Upper River Region, the Gambia, and its relationship with malaria infection in children. Malar J. 2018;17:1–14.

17. Padonou G, Sezonlin M, Gbedjissi G, Ayi I, Azondekon R DA, et al. Biology of Anopheles gambiae and insecticide resistance: Entomological study for a large scale of indoor residual spraying in south east Benin. J Parasitol Vector Biol. 2011;3:59–68.

18. Shcherbacheva A, Haario H, Killeen G. Modeling host-seeking behavior of African malaria vector mosquitoes in the presence of long-lasting insecticidal nets. Math Biosci. 2018;295:36–47.

19. Govella NJ, Chaki PP, Killeen GF. Entomological surveillance of behavioral resilience and resistance in residual malaria vector populations. Malar J. 2013:12:124.

20. Gillies MT, Coetzee M. A Supplement to the Anophelinae of the South of the Sahara (Afrotropical Region). South African Instit Med Res. 1987;55:1–143.

21. Scott JA, Brogdon WG, Collins FH. Identification of single specimens of the Anopheles gambiae complex by the polymerase chain reaction. Am J Trop Med Hyg. 1993;49:520–29

22. Fanello C, Santolamazza F, Della Torre A. Simultaneous identification of species and molecular forms of the Anopheles gambiae complex by PCR-RFLP. Med Vet Entomol. 2002; 16:461–4.

23. Bass C, Nikou D, Donnelly MJ, Williamson MS, Ranson H, Ball A, et al. Detection of knockdown resistance (kdr) mutations in Anopheles gambiae: A comparison of two new high-throughput assays with existing methods. Malar J. 2007;6:111.

24. Jones CM, Liyanapathirana M, Agossa FR, Weetman D, Ranson H, Donnelly MJ, et al. Footprints of positive selection associated with a mutation (N1575Y) in the voltage-gated sodium channel of Anopheles gambiae. Proc Natl Acad Sci. 2012;109:6614–19.

25. Martinez-Torres D, Chandre F, Williamson MS, Darriet F, Bergé JB, Devonshire AL, et al. Molecular characterization of pyrethroid knockdown resistance (kdr) in the major malaria vector Anopheles gambiae s.s. Insect Mol Biol. 1998; 7:179–84.

26. Ranson H, Jensen B, Vulule JM, Wang X, Hemingway J, Collins FH. Identification of a point mutation in the voltage-gated sodium channel gene of Kenyan Anopheles gambiae associated with resistance to DDT and pyrethroids. Insect Mol Biol. 2000;9:491–97.

27. Djogbenou LS, Weetman D, Dabire R, Ketoh G, Chandre F, Weill M, et al. The evolution of resistance to carbamates and organophosphate insecticides in Anopheles gambiae. Am J Trop Med Hyg. 2012.

28. Mitchell SN, Rigden DJ, Dowd AJ, Lu F, Wilding CS, Weetman D, et al. Metabolic and target-site mechanisms combine to confer strong DDT resistance in Anopheles gambiae. PLoS One. 2014;7:179–84.

29. Bass C, Nikou D, Vontas J, Williamson MS, Field LM. Development of high-throughput real-time PCR assays for the identification of insensitive acetylcholinesterase (ace-1R) in Anopheles gambiae. Pestic Biochem Physiol. 2010;96:80–5.

30. Bass C, Nikou D, Blagborough AM, Vontas J, Sinden RE, Williamson MS, et al. PCR-based detection of Plasmodium in Anopheles mosquitoes: A comparison of a new high-throughput assay with existing methods. Malar J. 2008;7:177.

31. Kent RJ, Thuma PE, Mharakurwa S, Norris DE. Seasonality, blood feeding behavior, and transmission of Plasmodium falciparum by Anopheles arabiensis after an extended drought in southern Zambia. Am J Trop Med Hyg. 2007;76(2):267–74.

32. Rebekah J. Kent and Douglas E. Norris. Am J Trop Med Hyg. Identif Mamm Blood Meals Mosquitoes By A Mult Polym Chain React Target Cytochrome B. 2005;73(2):336–342.

33. WHO. Guidelines for malaria vector control. 2019. Geneva: World Health Organization; 2019.

34. Silva APB, Santos JMM, Martins AJ, Tadei W, Thatcher B, Santos J, et al. Mutations in the voltage-gated sodium channel gene of anophelines and their association with resistance to pyrethroids – a review. Parasit Vectors. 2014;7:450.

35. Koukpo CZ, Fassinou AJYH, Ossè RA, Agossa FR, Sovi A, Sewadé WT, et al. The current distribution and characterization of the L1014F resistance allele of the kdr gene in three malaria vectors (Anopheles gambiae, Anopheles coluzzii, Anopheles arabiensis) in Benin (West Africa). Malar J. 2019;18:175.

36. Hamid-Adiamoh M, Amambua-Ngwa A, Nwakanma D, D’Alessandro U, Awandare GA, Afrane YA. Insecticide resistance in indoor and outdoor-resting Anopheles gambiae in Northern Ghana. Malar J. 2020;19:314.

37. Kabula B, Kisinza W, Tungu P, Ndege C, Batengana B, Kollo D, et al. Co-occurrence and distribution of East (L1014S) and West (L1014F) African knock-down resistance in Anopheles gambiae sensu lato population of Tanzania. Trop Med Int Heal. 2014;19:331–341.

38. Machani MG, Ochomo E, Amimo F, Kosgei J, Munga S, Zhou G, et al. Resting behavior of malaria vectors in highland and lowland sites of western Kenya: Implication on malaria vector control measures. PLoS One. 2020;15:e0224718.

39. Musiime AK, Smith DL, Kilama M, Rek J, Arinaitwe E, Nankabirwa JI, et al. Impact of vector control interventions on malaria transmission intensity, outdoor vector biting rates and Anopheles mosquito species composition in Tororo, Uganda. Malar J. 2019; 18(1):445.

40. Abong’o B, Gimnig JE, Torr SJ, Longman B, Omoke D, Muchoki M, et al. Impact of indoor residual spraying with pirimiphos-methyl (Actellic 300CS) on entomological indicators of transmission and malaria case burden in Migori County, western Kenya. Sci Rep. 2020;10(1):4518.

41. Machani MG, Ochomo E, Sang D, Bonizzoni M, Zhou G, Githeko AK, et al. Influence of blood meal and age of mosquitoes on susceptibility to pyrethroids in Anopheles gambiae from Western Kenya. Malar J. 2010:18:1–9.

42. Chaccour C, Killeen GF. Mind the gap: Residual malaria transmission, veterinary endectocides and livestock as targets for malaria vector control. Malar J. 2016;15:24.

43. Habtewold T, Prior A, Torr SJ, Gibson G. Could insecticide-treated cattle reduce Afrotropical malaria transmission? Effects of deltamethrin-treated Zebu on Anopheles arabiensis behavior and survival in Ethiopia. Med Vet Entomol. 2004;18:408–17.

44. Zoh DD, Yapi A, Adja MA, Guindo-Coulibaly N, Kpan DMS, Sagna AB, et al. Role of Anopheles gambiae s.s. and Anopheles coluzzii (Diptera: Culicidae) in Human Malaria Transmission in Rural Areas of Bouaké, in Côte d’Ivoire. J Med Entomol. 2020;57:1254–1261.

45. Killeen GF, Govella NJ, Lwetoijera DW, Okumu FO. Most outdoor malaria transmission by behaviorally-resistant Anopheles arabiensis is mediated by mosquitoes that have previously been inside houses. Malar J. 2016;15:1–10.

46. Bryan JH, Petrarca V, Di Deco MA, Coluzzi M. Adult behavior of members of the Anopheles gambiae complex in the Gambia with special reference to An. melas and its chromosomal variants. Parassitologia. 1987;29:221–49

47. Adamou A, Dao A, Timbine S, Kassogué Y, Yaro AS, Diallo M, et al. The contribution of aestivating mosquitoes to the persistence of Anopheles gambiae in the Sahel. Malar J. 2011;10:151.

48. Arcaz AC, Huestis DL, Dao A, Yaro AS, Diallo M, Andersen J, et al. Desiccation tolerance in Anopheles coluzzii?: the effects of spiracle size and cuticular hydrocarbons. J Exp Biol. 2016;219:1675–88

49. Durnez L, Coosemans M. Residual transmission of Malaria: An Old Issue for New Approaches. Anopheles mosquitoes - New insights into Malar vectors. 2013;671–704.

